# Behavioral and Neural Correlates of Temperament Traits: Insights from Temperament and Character Inventory and fMRI-Based Choice Tasks

**DOI:** 10.1101/2024.11.26.625354

**Authors:** Majid Abbasi Sisara, Seyed Hamid Khodadad Hosseini, Hossein Mahjoub, Asadollah Kordnaeij, Maryam Esmaeilinasab

**Affiliations:** Division of Biokinesiology and Physical Therapy, University of Southern California, Alcazar Street, 9006, Los Angeles, CA, USA; Department of Business Administration, Tarbiat Modares University, Tehran, Iran; Department of Psychology, Tarbiat Modares University, Tehran, Iran

**Keywords:** Cloninger Temperaments, Cognitive Dissonance, fMRI, free-choice paradigm

## Abstract

A cognitive conflict that is negatively arousing and results in a divergence in preference attitudes toward the chosen and rejected alternatives occurs when individuals are compelled to choose between alternatives that are similarly preferable. This phenomenon, which is frequently referred to as “cognitive dissonance,” is of interest in the fields of decision neuroscience and psychology. This study examines the behavioral and neural underpinnings of Cloninger’s temperament traits—Novelty Seeking (NS), Harm Avoidance (HA), Reward Dependence (RD), and Persistence (PS)—in a decision-making context.Functional magnetic resonance imaging (fMRI) and a modified free-choice paradigm were used to formalize the effect of Cloninger temperaments on cognitive conflict induced by choice. Behavioral analysis revealed significant individual differences across the four TCI dimensions, highlighting distinct variability in participants’ responses. Imaging data showed that participants with the highest and lowest scores in each temperament trait exhibited unique brain activation patterns during difficult and easy decision tasks. Notably, novelty-seeking was linked to heightened activation in brain regions associated with exploration and cognitive flexibility, while harm avoidance was associated with emotional processing and conflict detection. These findings provide deeper insights into how personality traits influence both behavior and neural responses during cognitive dissonance and decision-making tasks, offering implications for understanding individual differences in decision-related behaviors. The results of this study confirm the effect of temperaments on the degree of perceived dissonance of people, which can be used in many areas such as consumer behavior in marketing. The reason for this is the hesitancy of customers after purchasing the products.

**Significance Statement:** This study provides novel insights into how Cloninger’s temperament traits—Novelty Seeking, Harm Avoidance, Reward Dependence, and Persistence—modulate both neural and behavioral responses during decision-making processes. By combining fMRI with a modified free-choice paradigm, we reveal that temperament traits significantly impact cognitive dissonance, with distinct brain activation patterns corresponding to individual differences in decision conflict. These findings enhance our understanding of the neural basis of cognitive dissonance and have practical implications for areas such as consumer behavior, where temperament-driven decision conflict can influence post-purchase attitudes. This research highlights the importance of accounting for personality traits when examining decision-related behaviors and suggests that tailoring strategies based on temperament could improve decision outcomes in various applied fields.

---

Normative decision theory typically infers that our behaviors are indicative of our underlying preferences.Conversely, Festinger’s influential cognitive dissonance theory, which was introduced in 1957, contends that our actions are the ones that influence and modify our preferences (1). The mere act of selecting an item from two equally enticing alternatives has been shown to result in a change in an individual’s preferences toward the selected item, as evidenced by a multitude of studies (2).The term “cognitive dissonance” was first coined by Festinger to describe the distress we experience when contemplating psychologically incompatible ideas at the same time (3). We exert cognitive and behavioral effort to mitigate this dissonance and reestablish cognitive consistency in order to mitigate this cognitive dissonance (4). Cognitive dissonance is one of the most prominent contemporary models for comprehending the processing of inconsistent and conflicting cognitions due to the extensive empirical research and theoretical advancements of the field (5).

Researchers have found that when individuals face equally appealing options, they experience cognitive conflict, which is a state of mental discomfort (3). When confronted with challenging decisions, people exhibit longer reaction times and feel pressured to alleviate the discomfort caused by conflicting thoughts (2, 6). For instance, when someone faces conflicting thoughts (e.g., liking an item but rejecting it), they often adjust their attitude toward the options to reduce cognitive dissonance (3, 7).Cognitive dissonance is a term that psychologists frequently use to describe the phenomenon in which reduced cognitive conflict results in attitude adjustments.

The dorsal anterior cingulate cortex (dACC) has been identified as a potential neural correlate of cognitive conflict in recent neuroimaging findings (2, 6, 8–10). Van Veen et al. (8) utilized a paradigm requiring subjects to act against their own attitudes, revealing that such internal conflict is mirrored by dACC activation. Many researchers studied cognitive dissonance by using choices during fMRI between pairs of pre-rated DVD images, food items, music CDs, vacation destinations, and snack foods, respectively, as rated by the participants (2, 6, 9–12). During the trials, participants were presented with the option of selecting from options that were rated similarly (difficult) or contrastingly (simple). Although the included studies had different approaches to induce cognitive dissonance, they all reported some increased activity in the ACC or dACC during fMRI measurements.

The “free-choice paradigm” has been used in pioneering psychological research to demonstrate that individuals mitigate the cognitive tension of decision-making by perceiving chosen alternatives as more desirable and unchosen ones as less so (13). The free-choice paradigm involves a three-stage process where: first stage, Individuals express their initial preference for various options through subjective evaluation methods like rating or ranking. second stage, they are then presented with a choice between two of these options. Finally, they re-evaluate their preferences for each option postdecision. Research utilizing this paradigm has consistently found that after making a challenging decision between two appealing options, there is a noticeable shift in an individual’s preference. This shift is characterized by an increase in the attractiveness of the chosen option and/or a decrease in the attractiveness of the unchosen option. This phenomenon, known as the spreading of alternatives (SOA), is thought to lend support to the theory of cognitive dissonance (11, 14, 15). The SOA effect illustrates how decision-making can influence our perceptions and attitudes towards the options we consider (3).

Personality can be described as the collective traits that distinguish an individual (16, 17). This uniqueness depends on the interaction of genetic and environmental factors, and it is not surprising that every model and taxonomy that has attempted to describe the diversity of personality characteristics has considered this interrelationship (18). Cloninger’s psychobiological model stands out among these, delineating four fundamental temperaments—Novelty Seeking (NS); Harm Avoidance (HA); Reward Dependence (RD); and Persistence (P)—and three-character dimensions—Self-Directedness (SD); Cooperativeness (CO); and Self-Transcendence (ST), all quantifiable via the Temperament and Character Inventory (TCI) (19). The temperaments are typically tied to genetic factors, whereas the character dimensions are related to learned behaviors and cultural influences. It is posited that the balanced growth of these temperaments and characters is crucial for psychophysical well-being (20).

Personality theory (19) postulates that temperament dimensions are heritable traits in information processing and perceptual memory systems. Temperamental traits, which are primarily related to emotional responses and cognitive habits, are thought to be moderately heritable and relatively stable over life. Among the aspects of temperament, harm avoidance represents a tendency to worry pessimistically about the future and avoid uncertainty (19). people with a high degree of harm avoidance may see challenges as a source of anxiety and uncertainty. Novelty-seeking refers to the tendency to trigger and initiate novel discovery and response behavior (19). That is, individuals with a high degree of novelty prefer the new and therefore focus on or pay great attention to the new. Reward dependence represents the tendency to maintain and continue ongoing behavior and to depend on the approval of others (19). Maintaining and continuing an ongoing activity can improve proficiency in that activity over time. A high level of skill is likely to be trusted by others, and thus satisfy individuals with a high degree of reward dependence, motivating them to maintain and pursue the activity. Such a cycle of positive reinforcement gradually raises their skill level. Persistence is perseverance despite disappointment and fatigue (19). Individuals with high levels of persistence can stick with an activity despite frustration and fatigue. When they resolve deadlocks, their skill level increases markedly (21), helping them overcome difficult challenges. Table 1 describes individuals who scored high and low on four dimensions of temperament (22).

**Table 1.** A comparison of brain activation areas in subjects with the highest and lowest scores on the TCI Test based on different personalities traits. Based on the Temperament and Character Inventory (TCI) test, this table presents brain activation areas of subjects who scored highest and lowest in specific personality traits. The activated areas are correlated with subjects’ reported preference during Choice task (Figure1 B): hard condition vs. easy conditions. The hard condition is called self-difficult, and it is made up of trials that require participants to choose between two high-rated (or low-rated) items. A difficult condition occurs, for example, when participants are required to choose between two foods with ratings that are both higher than 7 (high-rated items). The easy conditions include: self-easy, system-difficult, and system-easy. The self-easy trials require participants to select between two items with different levels of preference, one of which is highly rated, while the other is low-rated. During a system-difficult trial, the computer selects between two items that participants rated both of them as high (or low) scores in preference task 1. During a system-easy trial, the computer selects between two items that participants rated one of them as high and the other one as low scores in preference task 1. In Table S1 to S7, you can find detailed information for all activation areas for each subject. A statistical threshold was set at *P <* 0.001 (FDR corrected) and the cluster size threshold is 5 voxels. In each panel, different traits are depicted along with corresponding brain activation areas. *(A)* Novelty Seeking This panel displays the brain areas that were activated during choice task in subjects with the highest (Subject09) and lowest (Subject04) scores in Novelty Seeking. In black, regions that were activated specifically for each subject are indicated, while in blue, regions that were activated for both subjects are indicated. Subject04 had more activated voxels in the Medial Frontal Gyrus and left Superior Parietal Lobule than Subject09, but Subject09 had more activated voxels in the left Superior Frontal Gyrus. *(B)* Harm Avoidance In this panel, the brain activation regions for participants with highest and lowest score in Harm Avoidance are illustrated. The highest scoring subject (Subject07) and the lowest scoring subject (Subject13) demonstrate distinct activation areas. The black lines indicate areas activated specifically for each subject, while the blue lines indicate areas activated for both subjects. There were more activated voxels in the Frontal Lobe, Limbic Lobe, Brodmann Area 32, Anterior Cingulate Cortex, and Medial Frontal Gyrus in Subject13 than in Subject07. *(C)* Reward Dependence This panel compares the activation regions of subjects with the highest (Subject05) and lowest (Subject01) scores in Reward Dependence. It was not found that there was a common region of activation between the two subjects during the choice task. In Subject01, which has the lowest reward dependency score, we can see that many brain regions were activated. In Subject05, we can see activation in five regions: left Superior Parietal Lobule, Precuneus, Parietal Lobe, Brodmann Area 7, and left Inferior Parietal Lobule. *(D)* Persistence The final panel shows the activated regions for Persistence, comparing the highest scoring subject (Subject11) with the lowest scoring subject (Subject07). There are black lines that indicate specific areas activated for each subject, and blue lines that indicate areas that are activated for both subjects. Subject11 had more activated voxels in the Frontal Lobe than Subject07, but Subject07 had more activated voxels in the Parietal Lobe.

| A) Novelty Seeking |  | B) Harm Avoidance |  |
| --- | --- | --- | --- |
| Highest score | Lowest score | Highest score | Lowest score |
| Subject09 | Subject04 | Subject07 | Subject13 |
| Hypothalamus | Precuneus | Inferior Parietal Lobule | Superior Frontal Gyrus |
| Amygdala | Fusiform Gyrus | Supramarginal Gyrus | Middle Frontal Gyrus |
| Insula | Brodmann Area 10 | Brodmann Area 40 | Superior Medial Frontal Gyrus (R) |
| Cingulate Gyrus | Cuneus | Brodmann Area 10 | Caudate (R) |
| Inferior Parietal Lobule | Brodmann Area 39 | Parietal Lobe | Cingulate Gyrus |
| Parietal Lobe | Supramarginal Gyrus (L) | Frontal Lobe | Frontal Lobe |
| Anterior Cingulate Cortex | Middle Temporal Gyrus (L) | Limbic Lobe | Limbic Lobe |
| Caudate | Medial Frontal Gyrus | Brodmann Area 32 | Brodmann Area 32 |
| Limbic Lobe | Superior Parietal Lobule (L) | Anterior Cingulate Cortex | Anterior Cingulate Cortex |
| Medial Frontal Gyrus | Superior Frontal Gyrus (L) | Medial Frontal Gyrus | Medial Frontal Gyrus |
| Superior Parietal Lobule (L) |  |  |  |
| Superior Frontal Gyrus (L) |  |  |  |
| C) Reward Dependence |  | D) Persistence |  |
| Highest score | Lowest score | Highest score | Lowest score |
| Subject05 | Subject01 | Subject11 | Subject07 |
| superior parietal lobule (L) | Inferior Orbital Frontal Cortex (L) | Precuneus (L) | Inferior Parietal Lobule |
| Precuneus | Inferior Frontal Gyrus | Cerebellum Posterior Lobe | Supramarginal Gyrus |
| Parietal Lobe | Brodmann Area 47 | Cerebellum (R) | Brodmann Area 40 |
| Brodmann Area 7 | Fusiform Gyrus | Cuneus | Brodmann Area 10 |
| Inferior Parietal Lobule (L) | Middle Frontal Gyrus | Parietal Lobe | Limbic Lobe |
|  | Brodmann Area 11 | Frontal Lobe | Brodmann Area 32 |
|  | Cerebellum Crus 1 (L) |  | Anterior Cingulate Cortex |
|  | Middle Orbital Frontal Gyrus (L) |  | Medial Frontal Gyrus |
|  | Uncus |  | Parietal Lobe |
|  | Parahippocampal Gyrus |  | Frontal Lobe |
|  | Insula |  |  |
|  | Limbic Lobe |  |  |
|  | Brodmann Area 28 |  |  |
|  | Inferior Temporal Gyrus |  |  |
|  | Brodmann Area 13 |  |  |
|  | Superior Temporal Gyrus |  |  |
|  | Amygdala |  |  |

With the development of brain imaging techniques, numerous neuroimaging studies have been conducted to investigate the neurobiological basis of human temperaments (23). Gardini et al. (24) demonstrated that traits such as novelty seeking (NS), harm avoidance (HA), reward dependence (RD), and persistence (PER) are heritable, stable over time, and dependent on genetics and biology. They examined whether structural variance in specific brain regions was reflected in individual differences in these personality traits. An MRI scan was performed on 85 participants who completed the Three-Dimensional Personality Questionnaire (TPQ). In order to analyze the correlation between individuals’ gray matter volume values and their personality trait scores, voxel-based correlation analyses were conducted. According to the results, structural variance in specific brain areas may explain individual differences in NS, HA, RD, and PER. In addition, Held-Poschardt et al. (25) conducted a study on patients with panic disorder (PD) using fMRI.

The results showed that the degree of activation in the ventral striatum during loss anticipation was positively correlated with harm avoidance and negatively correlated with novelty seeking. Markett et al. (26) utilized the harm avoidance scale from the Temperament and Character Inventory (TCI) as a trait marker to demonstrate increased functional connectivity within the insular salience of intrinsic connectivity networks (ICN). The results align with previous work, provide evidence for a potential biomarker of anxiety disorders, and most importantly, demonstrate a direct neural correlate of the personality trait harm avoidance in the absence of external stimulation. Furthermore, Jiang et al. (27) were motivated to realize the individualized prediction of four temperaments using whole-brain functional connectivity (FC). In predictions of the four temperament traits, brain connectivities that showed top contributing power were commonly concentrated in the hippocampus, prefrontal cortex, basal ganglia, amygdala, and cingulate gyrus. Finally, they demonstrated that a person’s temperament traits can be reliably predicted using functional connectivity strength within frontal-subcortical circuits, indicating that human social and behavioral performance can be characterized by a specific brain connectivity profile.

Many studies in the field of Neuroscience confirm the relationship between Cloninger’s temperament dimensions and the activity of different parts of the brain (17, 23, 28– 30). Investigations into the Cloninger personality model span both psychological and neuroscientific (17, 28, 29) disciplines. Moreover, the cognitive dissonance phenomenon is often examined through neuroscience research employing fMRI and the free-choice paradigm. This study is designed to explore how Cloninger’s temperament dimensions influence cognitive dissonance, utilizing the free-choice paradigm as its methodological approach. We demonstrated that different personalities have different brain activation patterns when they perform choice-tasks. It is important to note that we analysed fMRI data from individuals with high and low scores on each of Cloninger’s temperament dimensions, enabling us to compare the characteristics of each dimension with their brain regions that were active during choice-tasks.

## Results

### Behavioral Results

A detailed behavioral analysis was conducted based on the Temperament and Character Inventory (TCI) results. In the TCI assessment, four key dimensions were assessed: Novelty Seeking (NS), Harm Avoidance (HA), Reward Dependence (RD), and Persistence (PS). Each dimension was scored up to a maximum of 20 points. The box-plot in Figure 2 (A) illustrates the distribution of TCI 125 scores across all participants, highlighting the variability and central tendencies within each dimension. As can be seen in Figure 2 (B), the highest-scoring participants in each dimension are highlighted. The highest scores were achieved by Subject09 in NS, Subject07 in HA, Subject05 in RD, and Subject11 in PS. On the other hand, Figure 2 (C) highlights participants with the lowest scores in each dimension. There were four subjects with the lowest scores in each dimension:

Subject04 in NS, Subject13 in HA, Subject01 in RD, and Subject07 in PS. Individual differences in temperament and personality traits are evident in these results, providing valuable insights for future psychological and behavioral studies. These subjects were chosen to undergo MRI brain scanning during a choice task. When we use participants from both ends of a dimension, we can observe differences in brain activation.

The experimental tasks results showed a significant decrease in preference for rejected foods in the PostEx-Choice condition (P *<* 0.01) when compared with rejected foods in the SelfEasy condition (P *<* 0.01) in Figure 3. As suggested by Chen and Risen (9), a typical free-choice paradigm produces preference changes even without cognitive dissonance. In the SelfDifficult condition, preferences for rejected foods dropped even further, but the change was significantly less than preference changes for rejected items observed in the PostEx-Choice condition (P *<* 0.05) in Figure 3, providing evidence of choice-induced preference changes. For the “Selected” category, preferences change negatively for all conditions, with self-difficulty, self-easy, and computer ratings decreasing by -0.10, -0.30, and -0.31, respectively, and the post-ex choice rating decreasing by -0.18. No significant differences were detected between them for selected foods.

### Imaging Results

Among subjects with differing scores on the Temperament and Character Inventory (TCI), the fMRI data indicated significant differences in brain activation pattern. A comparison of brain activity during the Choice task (Figure1 B) between subjects with the highest and lowest scores was made for each of the four personality traits: novelty seeking, harm avoidance, reward dependence, and persistence. We first analyzed Choice Task scans for each subject using general linear models (GLMs) (see Methods for details about GLMs) and found brain areas that were activated when in difficult vs easy conditions. The hard condition is called self-difficult, and it is made up of trials that require participants to choose between two high-rated items. A difficult condition occurs, for example, when participants are required to choose between two foods with ratings that are both higher than 7 (high-rated items). The easy conditions include: self-easy, system-difficult, and system-easy. The self-easy trials require participants to select between two items with different levels of preference, one of which is highly rated, while the other is low-rated. During a system-difficult trial, the computer selects between two items that participants rated both of them as high scores in preference task 1 (Figure1 A). During a system-easy trial, the computer selects between two items that participants rated one of them as high and the other one as low scores in preference task 1. The following sections describe these differences in detail.

**Fig. 1.**
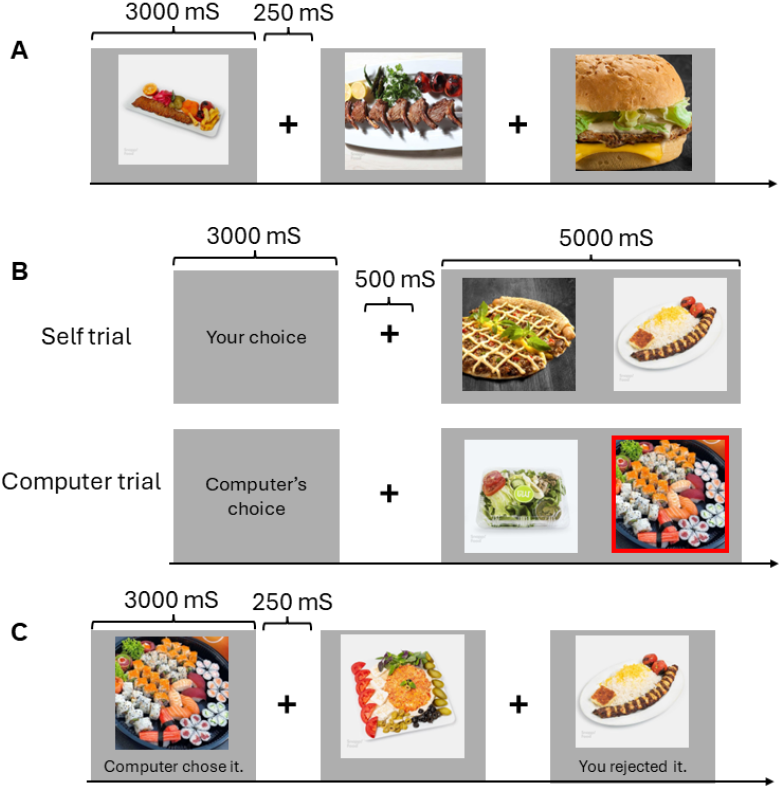
Experimental tasks. (***A***) **Preference task 1**. On a four-point scale, subjects rated their preference for each food item after seeing it for 3000 ms. (***B***) **Choice task**. An example of self-trial and computer-trial from the Choice task. During a self-trial, participants selected one of two foods based on their preferences. For the computer trial, participants had to identify the item randomly selected by the computer (highlighted by a red square). Cues were presented for 3000 ms, followed by a brief delay of 500 ms, and then two foods were presented for 5000 ms. (***C***) **Preference task 2**. The subjects were presented with the same food stimuli as in Preference task 1, and they were once again asked to rate their preference. Additionally, under each picture, past decisions by participants or a computer were displayed (e.g., “You rejected it”, etc.). Only images of foods were presented in the absence of items utilized for the Choice task.

#### A. Novelty Seeking

Subjects with high Novelty Seeking scores demonstrated distinct neural activation compared to those with low scores. The brain areas that were activated in Subject09, who scored the highest, are the following: Hypothalamus, Amygdala, Insula, Cingulate Gyrus, Inferior Parietal Lobe and Parietal Lobe, anterior Cingulate Cortex, Caudate, and Limbic Lobe. However, Subject04, the lowest-scoring subject, activated the following brain areas: Precuneus, Fusiform Gyrus, Brodmann Area 10, Cuneus, Brodmann Area 39, left Supramarginal Gyrus, and left Middle Temporal Gyrus. Specifically, Subject09, exhibited greater activation in the left Superior Frontal Gyrus and Precuneus, regions associated with exploratory behavior and decision-making under uncertainty (31, 32). In contrast, Subject04, had more prominent activation in the Medial Frontal Gyrus and left Superior Parietal Lobule, which are linked to self-referential thinking and attentional control (33, 34). Those who exhibit a higher level of novelty-seeking may engage brain regions associated with exploration and cognitive flexibility, while those who achieve lower scores tend to rely more heavily on brain regions related to self-monitoring and control.

#### B. Harm Avoidance

In the Harm Avoidance trait, Subject07 (highest score) and Subject13 (lowest score) displayed notable differences in brain activation. Subject07 showed activation in the following brain regions: Inferior Parietal Lobule, Supramarginal Gyrus, Brodmann Area 40 and 10, and Parietal Lobe. According to Subject13, superior frontal, middle frontal, right superior medial frontal, right caudate, and right cingulate gyrus were activated. The highest scorer showed increased activation in the Inferior Parietal Lobule, and Caudate, areas involved in emotional processing and avoidance behavior (35). Conversely, the lowest scorer activated the Limbic Lobe, Anterior Cingulate Cortex, and Medial Frontal Gyrus, regions associated with conflict detection and emotional regulation (36, 37). High harm avoidance may result in heightened activation in areas associated with anxiety and risk aversion, while low harm avoidance may involve circuits associated with regulating emotional responses during decision-making.

#### C. Reward Dependence

The comparison between high and low Reward Dependence scores revealed distinct patterns of brain activity. Subject05, who had the highest score, showed activation primarily in the left Superior Parietal Lobule, Precuneus, left Inferior Parietal Lobule, and Brodmann Area 7, areas associated with the integration of sensory information and visuospatial attention (33, 38). Subject01, who scored the lowest, had widespread activation across regions such as the left Inferior Orbital Frontal Cortex, Inferior Frontal Gyrus, Brodmann Area 11, 13, 28, 47, left Cerebellum Crus 1, Middle Frontal Gyrus, left Middle Orbital Frontal Gyrus, Uncus, Insula, Limbic Lobe, Inferior Temporal Gyrus, Superior Temporal Gyrus, Amygdala, and Parahippocampal Gyrus, which are implicated in decision-making, cognitive control, and memory processing (39–41). As a result of differential engagement in these areas, individuals with higher reward dependence are likely to be more focused on sensory processing and attention management, while those with lower scores tend to engage broader cognitive control and memory networks during decision making.

#### D. Persistence

In the Persistence trait, the highest scorer (Subject11) showed increased activation in the left Precuneus, Frontal Lobe, right Cerebellum, Cuneus, and Cerebellum Posterior lobe regions involved in executive functions and goal-directed behavior (42, 43). The lowest scorer (Subject07) displayed greater activation in the Parietal Lobe and Inferior Temporal Gyrus, associated with visuospatial processing and attentional shifts (44, 45). According to this pattern, high persistence appears to be associated with an increase in executive control, while low persistence may be associated with an increase in attentional reorientation and visual processing.

Across all traits, the data indicate that different personality characteristics are associated with distinct neural activation patterns during decision-making tasks. According to these findings, personality traits are associated with specific brain regions. For example, high novelty-seeking is often correlated with increased activity in the prefrontal cortex and parietal regions, supporting their role in adaptive decision-making (19, 46). Similarly, harm avoidance has been associated with heightened activation in limbic and prefrontal regions, reflecting its connection to anxiety and risk assessment (47).

## Discussion

According to Cloninger (2004) (22), individuals scoring high and low on the four temperament dimensions are: **A) Novelty Seeking:** High scorers tend to be exploratory, impulsive, extravagant, and irritable, whereas low scorers are more reserved, rigid, frugal, and stoic. **B) Harm Avoidance:** High scores are linked to traits like pessimism, fearfulness, shyness, and fatigability, while low scores align with optimism, daringness, outgoingness, and vigor. **C) Reward Dependence:** Those scoring high tend to be sentimental, sociable, warm, and sympathetic, while low scores indicate a critical, aloof, detached, and stoic nature. **D) Persistence:** High scores show eagerness, determination, ambition, and perfectionism, while low scores reflect apathy, being spoiled, underachievement, and pragmatism (Appendix SI Table S1).

A comparison of the brain regions activated by high and low Novelty Seeking (NS) subjects reveals distinct patterns that are consistent with their known functional attributes.

### High Novelty Seeking

In high NS scorers, the Amygdala, Hypothalamus, Insula, and Anterior Cingulate Cortex (ACC) are activated, indicating an emphasis on emotion, reward processing, and risk assessment. As well as facilitating decision-making and emotional responses to risk and reward, these regions, especially the Amygdala and Insula, also play an important role in the appraisal of new or rewarding stimuli (48, 49). Hypothalamus and ACC play major roles in modulating responses to reward-based stimuli, especially reward sensitivity and behavior regulation (50). A high NS individual likely exhibits exploratory behavior due to the integration of emotional processing with action planning in the Limbic Lobe, including the Caudate (51). Parietal lobe and superior parietal lobule activity may also reflect attention and spatial processing, which facilitates exploration and interactions with new environments. Motivation, conflict monitoring, and decision-making under uncertainty are supported by the Medial Frontal Gyrus and Cingulate Gyrus. They are associated with a drive for new experiences (52, 53).

### Low Novelty Seeking

Subjects with low novelty seeking exhibit activation in regions associated with self-referential thinking, social cognition, and visual processing, such as the Precuneus, Fusiform Gyrus, and Brodmann Areas 10 and 39. This region of the brain is associated with episodic memory and self-reflection, and may indicate that these individuals are more reserved or cautious when making choices (54). The Fusiform Gyrus assists in processing detailed visual information, possibly aligning with a more analytical approach to new stimuli. Additionally, supramarginal and middle temporal gyrus activations contribute to social perception and emotional regulation, and these processes appear to be crucial for restrained, calculated behavior often observed in lower NS individuals (55). Both high and low NS scorers have activations in the Medial Frontal Gyrus and Superior Parietal Lobule, but they may contribute differently to novelty processing, reflecting differences in exploratory behavior and self-regulation (56, 57). According to these findings, high NS individuals are neurologically predisposed to react to novel stimuli in a combination of an emotional approach and a reward-oriented approach, while low NS individuals use brain areas more focused on reflecting, cautioning, and being socially aware.

### High Harm Avoidance

Individuals with high Harm Avoidance will exhibit activation in areas like the Inferior Parietal Lobule, Supermarginal Gyrus, Brodmann Area 40, and Brodmann Area 10 which are involved in cautious, riskaverse behavior. Supramarginal Gyrus and Inferior Parietal Lobule play roles in social cognition, and they are typically more prominent in harm-avoidant people. HA individuals are more inclined toward caution and heightened sensitivity to feedback when they have the supramarginal gyrus, which is involved in interpreting social cues and maintaining attention on potential risks (58, 59). A crucial part of managing conflict and regulating emotions is the Brodmann Area 32 and Anterior Cingulate Cortex (ACC). In order to avoid harm, these regions monitor and react to potential risks in the environment. According to studies, the ACC promotes readiness to avoid threats, aligning with the characteristics of HA (60, 61). Individuals with HA often maintain caution and avoid negative outcomes due to the Limbic Lobe, Medial Frontal Gyrus, and Frontal Lobe, which are associated with processing emotions, self-regulation, and executive control. In decision-making, the Medial Frontal Gyrus plays an important role, since it supports reflective processing under uncertainty, likely contributing to harm-avoidance (62, 63). A high HA individual is likely to benefit from Brodmann Area 10, because it focuses on strategic planning and long-term evaluation. Negative outcomes frequently activate this region when decision-making is framed around preventing them (64).

### Low Harm Avoidance

Individuals with low Harm Avoidance scores were shown to have activation of regions such as the Superior Frontal Gyrus, Middle Frontal Gyrus, Superior Medial Frontal Gyrus (R), Caudate (R), and Cingulate Gyrus, suggesting that the neural framework enables a more flexible, exploratory approach to decisionmaking than a cautious, risk-averse approach. An adaptable approach to decision-making is supported by the Superior and Middle Frontal Gyri, which are involved in self-regulation, cognitive flexibility, and working memory. Specifically, the Superior Medial Frontal Gyrus (R) facilitates self-referential thinking, allowing individuals with low HA to be more flexible and exploratory (55, 65). The Caudate (R) facilitates more routine behavior and less risk-focused behavior by facilitating habit formation and action selection. A low HA individual is less likely to show high caution, favoring established behaviors and decisions that do not prioritize harm avoidance (66). Activation of the Brodmann Area 32 and Anterior Cingulate Cortex occurs in both high and low HA individuals, although the roles they play may be distinct. The activation of the ACC may enable exploratory choices rather than avoidance-focused processing in individuals with low HA (67). Individuals with high and low HA levels activate the Limbic Lobe and Medial Frontal Gyrus, but they may function differently. A low HA individual’s emotional processing is more balanced than a high HA individual’s, which suggests that a lower HA individual experiences reward and risk equally (68). Using this distinct pattern of brain activation, it is evident that high HA individuals use the brain regions related to vigilance, social processing, and emotional regulation, while low HA individuals have a neural signature related to flexibility, exploration, and reduced sensitivity to risks.

### High Reward Dependence

The regions of the brain active in individuals with a high reward dependency include the Superior Parietal Lobe (L), Precuneus, Parietal Lobe, Brodmann Area 7, and Inferior Parietal Lobule (L), which indicate an increased focus on attention, social awareness, and reward-related behavior. Social cognition, spatial awareness, and self-referential thought are all key functions of the Superior Parietal Lobule (L) and Precuneus, which align with high RD individuals’ tendency to seek social approval and rely on social cues to make decisions (69, 70). As the Precuneus is associated with self-reflection and empathy, it plays a key role in supporting behaviors based on social dependency and empathy. A significant neural involvement in attentional processes and visual-spatial processing is evident in the Parietal Lobe and Brodmann Area 7. These regions are essential for orienting attention, likely aiding high RD individuals in remaining sensitive to reward-related cues, especially those that are socially oriented (71). For people with high RD scores who depend on positive reinforcement from others, the inferior parietal lobule (L) is crucial to understanding social context and intention. Reward dependency aligns with this region’s role in processing social cues, since high RD individuals value rewards associated with social interactions(72).

### Low Reward Dependence

A neural framework that favors independent processing, emotional detachment, and self-sufficiency is evident in individuals with low reward dependence, including the Inferior Orbital Frontal Cortex (L), Inferior Frontal Gyrus, Brodmann Area 47, Fusiform Gyrus, and Middle Frontal Gyrus. There is an association between emotional regulation and inhibitory control in the Inferior Orbital Frontal Cortex (L), Inferior Frontal Gyrus, and Brodmann Area 47. Individuals with low RD can detach from social reinforcement and process choices based on internal motivations rather than social rewards(73, 74). A tendency to prioritize internal goals over external rewards can also be reflected in the Middle Frontal Gyrus due to its activation, which also supports cognitive control and self-reliance in decision-making. A key function of the Fusiform Gyrus is to process details of visual information, particularly to recognize faces and other socially relevant stimuli. The activation of this pathway in individuals with low RD may indicate a preference for autonomy over social dependence (75). There is a connection between emotion processing and memory in the Cerebellum Crus I (L), Middle Orbital Frontal Gyrus (L), and Uncus (L). These areas foster emotional resilience in low RD individuals, enabling them to make decisions with minimal external rewards (76). For emotion processing, empathy, and social perception, other regions, including the Parahippocampal Gyrus, Insula, Amygdala, Superior Temporal Gyrus, and Brodmann Area 13, are important. Activation in these areas may, however, indicate that individuals with low RD are capable of processing social information without strong emotional or reward reinforcement, supporting independent and self-regulated decision-making (77, 78). As a result of this pattern of neural activity, it appears that high RD individuals rely heavily on areas that are associated with social perception, empathy, and attentional processing, while low RD individuals rely heavily on regions related to self-sufficiency, emotional regulation, and cognitive control.

### High Persistence

Brain activations in high persistence individuals include the Precuneus (L), Cerebellum Posterior Lobe, Cerebellum (R), Cuneus, Parietal Lobe, and Frontal Lobe. Sustained attention, goal-directed behavior, and mental resilience are associated with these regions. Precuneus (L) plays a crucial role in self-referential processing and maintaining focus on long-term goals, which is related to high persistence individuals’ efforts and commitment to long-term objectives(79). This region supports continuous attention and internal motivation, essential for individuals with high levels of persistence. Motor control and cognitive tasks that require precision and planning are performed in the Cerebellum Posterior Lobe and Cerebellum (R). Specifically, the cerebellum aids individuals who maintain consistent efforts to reach their goals by adjusting behavior based on feedback(80, 81). A high level of Cuneus activation may be linked to visual processing and spatial orientation in high persistence individuals. It may enhance the ability to process environmental cues, which may be useful when performing goal-oriented tasks that require sustained visual attention and decision-making (82). A key component of executive function is the Parietal Lobe and Frontal Lobe, which help us plan, make decisions, and maintain focus for long periods of time. By regulating goal-directed behavior, high Persistence individuals manage and overcome obstacles (83).

### Low Persistence

Anterior Cingulate Cortex (ACC), Medial Frontal Gyrus, Parietal Lobe, and Frontal Lobe are active in low persistence individuals. The inferolateral lobule, supramarginal gyrus, limbic lobe, and Brodmann area 32 are also active. The neural profile of these regions may support flexible decision-making, emotional responsiveness, and lower resilience to sustained effort. It is believed that the superior parietal lobule and supramarginal gyrus are involved in processing social and emotional stimuli, which may contribute to the lower levels of persistence in people by focusing on immediate social rewards rather than longterm goals (84, 85). In these areas, individuals with low persistence might adjust their efforts in response to social feedback or emotional reactions rather than goal endurance. In individuals with low persistence, Brodmann Area 40 and Brodmann Area 10 may reflect a focus on present-oriented goals rather than future-oriented ones. For example, Brodmann Area 10 is linked to decision-making and strategic flexibility rather than sustained effort (86). Emotional processing and conflict monitoring are associated with the Limbic Lobe, Brodmann Area 32, and ACC. There is evidence that people with low persistence may be more sensitive to emotional or motivational fluctuations, affecting their ability to remain consistent in goal-directed activities (87). Medial Frontal Gyrus, Parietal Lobe, and Frontal Lobe are also activated, reflecting cognitive functions associated with task engagement. However, low-persistent individuals may employ adaptive disengagement and task switching instead of prolonged focus, thus resulting in lower perseverance (88). Based on this activation pattern, individuals with high Persistence engage brain regions associated with sustained effort, self-regulation, and executive function, while those with low Persistence utilize areas associated with adaptive flexibility, social and emotional responsiveness, and present-oriented processing. Understanding the neural profiles associated with temperament dimensions such as novelty seeking (NS), harm avoidance (HA), reward dependency (RD), and persistence (P) can greatly enhance marketing strategies by enabling personalized approaches. High NS individuals are drawn to new experiences and adventure, making them ideal targets for innovative product launches and marketing campaigns focused on novelty and excitement. Conversely, low NS individuals may respond better to marketing that emphasizes reliability, security, and practicality. Tailoring content to highlight long-term value can appeal to those who prefer more calculated decision-making.

Similarly, a nuanced understanding of HA and RD can guide more effective marketing. High HA individuals, who are more risk-averse, respond well to reassurance and messages that minimize negative outcomes, with products promoting safety and comfort being particularly appealing. Low HA individuals, on the other hand, are more exploratory and flexible, responding well to messaging that emphasizes freedom and spontaneity. High RD individuals, who value social connections and external validation, are best targeted with strategies that include social proof, such as celebrity endorsements or community engagement. Low RD individuals, who prioritize self-sufficiency, are more responsive to marketing that emphasizes autonomy and personal achievement. The neural profiles associated with persistence also offer marketing insights. High persistence individuals, who are goal-oriented and resilient, are likely to engage with campaigns that highlight long-term progress and success, such as those focused on career advancement or fitness. Marketers can use messaging that focuses on milestones and rewards for sustained effort. In contrast, low persistence individuals, who are more flexible and seek immediate rewards, respond better to campaigns offering quick benefits, instant gratification, or limited-time promotions.

Understanding how different brain regions influence behavior further refines strategies for high and low RD groups. High RD individuals, who seek social validation, respond well to influencer partnerships and community-based marketing. Low RD individuals, who value autonomy, are drawn to messages that emphasize self-reliance and personal goals. By tailoring marketing strategies to these neural profiles, brands can create more personalized, relevant experiences that foster stronger customer engagement, loyalty, and conversion rates. This approach allows marketers to align campaigns with the psychological drivers of their target audiences.

## Methods

### Subjects

Over 234 participants took the Temperament and Character Inventory (TCI) test (52% women, 48% men; 18-65 years old) (see Appendix SI Table S9). To obtain MRI images, 14 male participants (age range 30-40) were selected. In this paper, We processed choice task fMRI data of seven of these participants who scored highest or lowest in each of the four personality traits. None of the participants had a history of neurological or psychiatric illness. Informed consent was obtained from all subjects, and the study was approved by the Research Ethics Committee of Iran University of Medical Sciences (Approval ID: **IR.IUMS.REC.1400.928**).

### Tasks and Experimental Protocol

The present investigation was divided into four separate parts: (A) Preference task 1, (B) Choice task, (C) Preference task 2, and (D) PostExperimental Choice task. Unless the Post-Experimental Choice task was conducted in the fMRI scanner, all other tasks were performed there (2).

The Preference task 1 (Fig.1A) was a questionnaire that asked the participants to rate their preference for a food that was displayed on the screen on a scale of 1-8. A score of 1 indicates that they did not like the food at all, while a score of 8 indicated that they disliked it exceedingly. The Choice task consisted of two distinct types of trials: the Self trial and the Computer trial (Fig.1B). Self-trial subjects were instructed to select one product from two that were simultaneously displayed on a screen (Fig.1B, Upper). Subjects conducted the computer trial (Fig.1B, Lower) by clicking on the button that corresponded to a randomly selected food (highlighted by a red square). The same 160 foods were presented on the screen in Preference Task 2, and the subjects were asked to rate their preferences. One significant distinction was that the past decisions of subjects (or computers) during the Choice task were displayed under each of the foods included in the pairs presented during the Choice task (e.g., “You chose it,” “You rejected it,” “Computer chose it,” or “Computer rejected it”) (Fig.1C). The image of a stimuli that was not utilized in the Choice task was displayed exclusively on the screen, as was the case in Preference task 1.

**Fig. 2.**
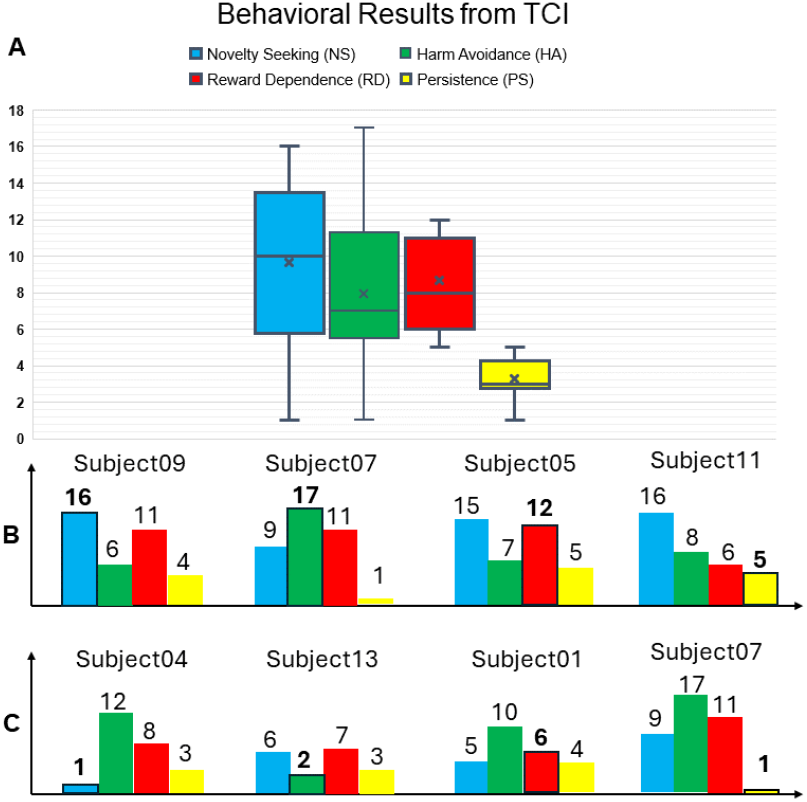
Behavioral result of the Temperament and Character Inventory (TCI). (***A***) The box-plot of TCI 125 results over all participants. It includes four dimensions: Novelty Seeking (NS), Harm Avoidance (HA), Reward Dependence (RD), and Persistence (PS). Grades can be given on each dimension up to 20. (***B***) Participants with the highest grade in each dimension. For example, among all participants, Subject09 has the highest score (16 out of 20) in Novelty Seeking. (***C***) Participants with the lowest grade in each dimension. For example, Subject04 has the lowest score (1 of 20) in Novelty Seeking among all participants.

**Fig. 3.**
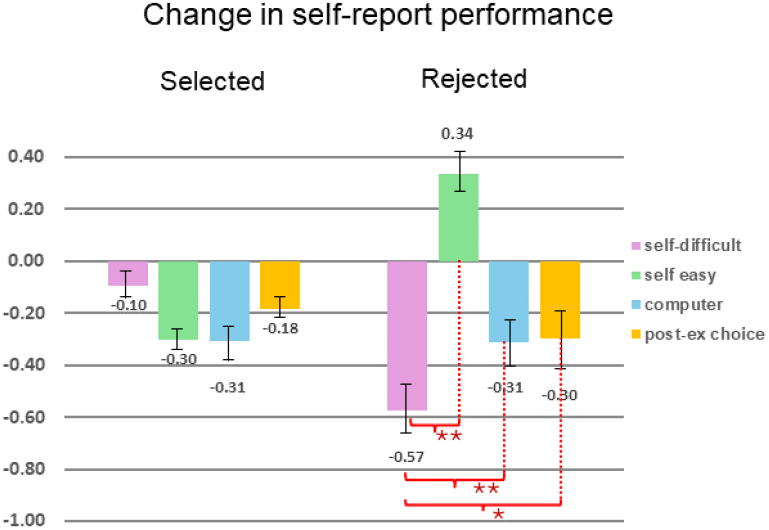
The behavioral result of the preference change. The bars in each condition indicate the difference in preference ratings from Preference task 1 to Preference task 2 (preference ratings in Preference task 2 - preferences in Preference task 1). *∗* P *<* 0.05; *∗∗* P *<* 0.01 (paired t-test, one-tailed). The error bars indicate the standard error of the mean (SEM).

**Fig. 4.**
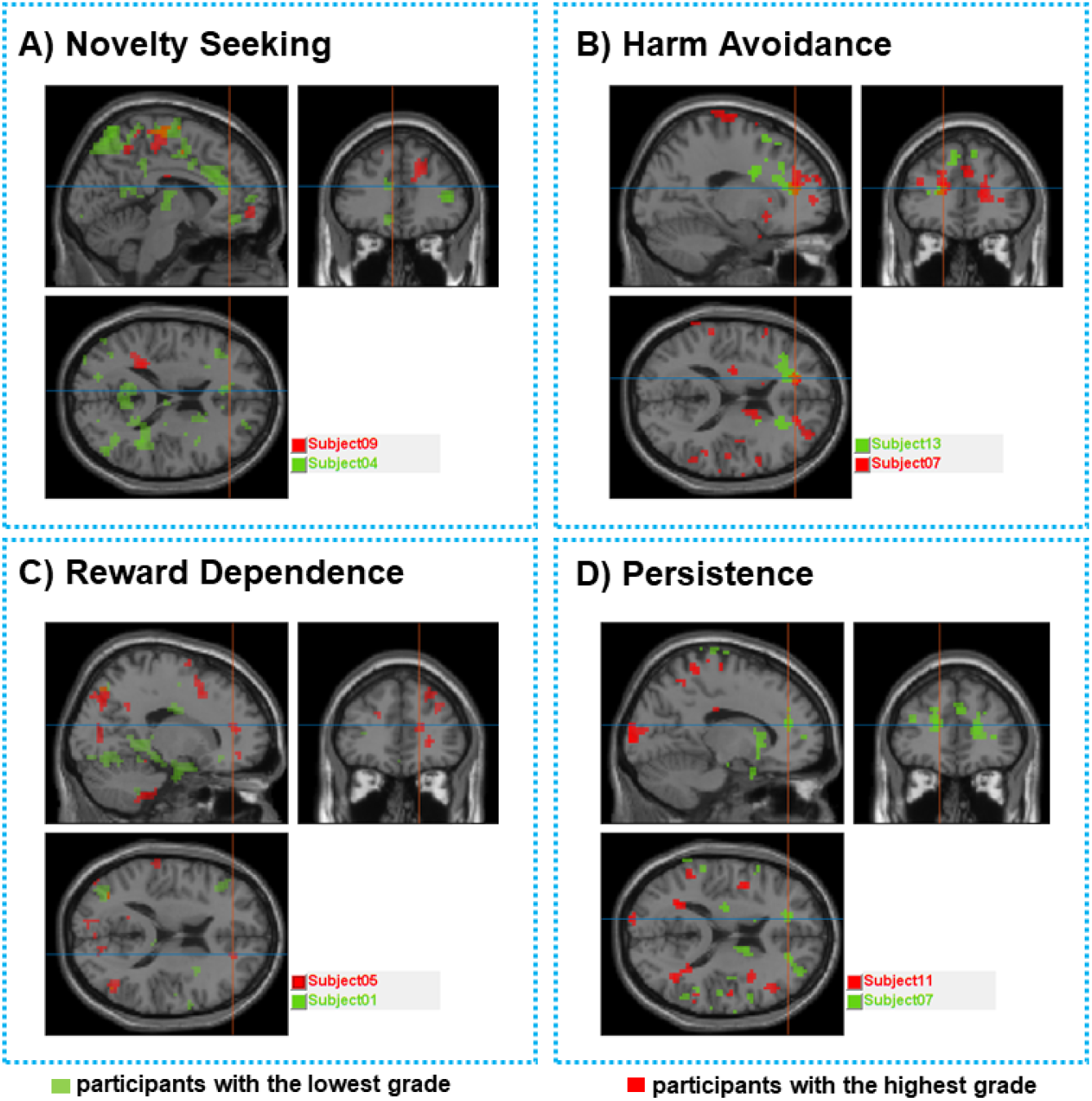
Brain activation maps of choice task from subjects with highest and lowest scores in different personality traits from the TCI Test. This figure presents the brain activation maps of subjects who scored highest (in red) and lowest (in green) in various personality traits as measured by the Temperament and Character Inventory (TCI) test. Areas correlated with subjects’ reported preference during Choice task (Figure1 B) Almost the entire cingulate cortex (from anterior to posterior) was activated. All activations are listed in Table S1. A statistical threshold was set at *P* < 0.001 for height (uncorrected) and cluster *P* < 0.05 (corrected for multiple comparisons). Each trait is depicted in a separate panel with corresponding brain images in different orientations to provide comprehensive views of the activation regions. ***(A)* Novelty Seeking** This panel displays the brain activation patterns of subjects with the highest (Subject09, red) and lowest (Subject04, green) scores in Novelty Seeking. The images show the sagittal, coronal, and axial views, highlighting the differential activation in brain regions associated with novelty and exploration behaviors. ***(B)* Harm Avoidance** In this panel, the brain activation maps for Harm Avoidance are illustrated. The highest scoring subject (Subject07, red) and the lowest scoring subject (Subject13, green) demonstrate distinct activation patterns. The images include sagittal, coronal, and axial views, focusing on areas linked to anxiety and fear-related responses. ***(C)* Reward Dependence** This panel compares the activation maps of subjects with the highest (Subject05, red) and lowest (Subject01, green) scores in Reward Dependence. The different views (sagittal, coronal, and axial) emphasize the brain regions involved in reward processing and social attachment. ***(D)* Persistence** The final panel shows the activation maps for Persistence, comparing the highest scoring subject (Subject11, red) with the lowest scoring subject (Subject07, green). The views provided highlight the brain regions associated with perseverance and sustained effort.

The PostEx-Choice task was conducted outside of the fMRI scanner following the completion of Preference task 2. Subjects were requested to select their preferred food from a pair of options presented on the computer trial of the Choice task. Afterward, the subjects evaluated their familiarity with 160 foods using an eight-point scale.

### Functional MRI Data Collection and Analysis

MRI scanning was conducted using a 3.0-Tesla scanner (MAGNETOM Prisma, Siemens, Germany) that was equipped with a 20channel head coil array for signal reception. Functional imaging was conducted using a T2*-weighted echo planar imaging (EPI) sequence that was sensitive to BOLD contrast. The sequence was configured with the following parameters: a repetition time (TR) of 3000 ms, an echo time (TE) of 30 ms, a flip angle of 90°, a 72 × 64 acquisition matrix, a field of view (FOV) of 240 mm, an in-plane resolution of 3 × 3 mm, and 36 axial slices with a slice thickness of 3 mm and an interslice gap of 0.5 mm. T1-weighted, magnetization-prepared rapidacquisition gradient echo (MP-RAGE) pulse sequences were employed to acquire high-resolution structural images with a spatial resolution of 1×1×1mm.. The subject’s cranium was confined by a firm padding, which impeded its ability to move. Visual stimuli were displayed outside the scanner chamber using the screen mounted on the head coil. The subject’s responses were collected using magnet-compatible response receptacles. Three consecutive runs were conducted to acquire the EPI images. The initial four scans of each session were discarded in order to account for T1 equilibration effects.

The data were analyzed using SPM12 (Wellcome Department of Imaging Neuroscience, London, UK). Each volume acquired from each subject was realigned to account for minor movements between scans. This process generated a mean image and an aligned set of images for each subject. Subsequently, we adjusted the realigned images to account for variations in slice acquisition times. The mean of the realigned EPI images was used to coregister the T1-weighted structural MRI images of each participant, which were then segmented to separate the gray matter. Gray matter was normalized using a template image that was derived from the Montreal Neurological Institute (MNI) reference brain. Voxels were resampled at a resolution of 2 mm × 2 mm × 2 mm. A Gaussian kernel of 8 mm full-width and halfmaximum was employed to flatten the EPI images after they were normalized to the MNI template.

At the single subject level, we applied whole-brain general linear model (GLM) analysis as an event-related model. A canonical hemodynamic response function was convolutioned with the hemodynamic response to stimulus onset for each participant. In this paper we only analyzed the choice task fMRI data since the primary objective was to investigate whether different personalities have different brain activations during cognitive dissonance or when their preferences change as a result of their choice. There were four experimental conditions: (1) the Self-Easy condition, (2) the Self-Difficult condition, (3) the Computer-Easy condition, and (4) the Computer-Difficult condition. In difficult conditions, there are two food items presented on the screen with high scores. In order to determine which brain region was activated during the choice task, we used the following contrast in a

t-test:

*contrast* = (*self* -*Dif f icult*) *− others* and others are all other conditions:

*other* = (*self* -*Easy*) + (*computer*-*Dif f icult*) + (*computer*-*Easy*)

Using a voxel-by-voxel t-test and a p-value threshold (*p <* 0.05) the Table.S2 to Table.S8 were calculated.

### Data Archival

You can find all the participants’ fMRI data (dicom format) on the link below.

## ACKNOWLEDGMENTS

The authors would like to acknowledge the Iranian National Brain Mapping Laboratory (NBML), Tehran, Iran, for providing the data acquisition service for this research work.

## Author contributions

S.H.K.H., A.K., M.E. and H.M. designed research; S.H.K.H., M.A.S., and H.M. performed research; S.H.K.H., M.A.S., H.M., A.K. and., M.E. contributed new reagents/analytic tools; M.A.S. and H.M. analyzed data; and M.A.S., S.H.K.H. and H.M. wrote the paper.

## Supporting Information

**Table S1.** Descriptors of individuals who score Hight and Low on the Four Temperament Dimensions.

| Temperament Dimension | Descriptors of Extreme Variants |  |
| --- | --- | --- |
|  | High | Low |
| **Harm Avoidance** | • Pessimistic<br>• Fearful<br>• Shy<br>• Fatigable | • Optimistic<br>• Daring<br>• Outgoing<br>• Vigorous |
| **Novelty Seeking** | • Exploratory<br>• Impulsive<br>• Extravagant<br>• Irritable | • Reserved<br>• Rigid<br>• Frugal<br>• Stoic |
| **Reward Dependence** | • Sentimental<br>• Sociable<br>• Warm<br>• Sympathetic | • Critical<br>• Aloof<br>• Detached<br>• Stoic |
| **Persistence** | • Eager<br>• Determined<br>• Ambitious<br>• Perfectionist | • Apathetic<br>• Spoiled<br>• Underachiever<br>• Pragmatist |

**Table S2.** The number of voxels in each cluster that are activated during the choice task in subject01. Sabject01 had the lowest score in Reward Dependence of TCI test. The contrast was defined as (self-difficult - other conditions) and the p-value threshold was set at (P<0.05).

| Number of voxels | ROIs |
| --- | --- |
| 42 | Inferior Frontal Gyrus |
| 27 | Occipital Lobe |
| 26 | brodmann area 47 |
| 18 | Occipital_Inf_L (aal) |
| 17 | Middle Occipital Gyrus |
| 14 | Fusiform_L (aal) |
| 13 | Fusiform Gyrus |
| 13 | Fusiform Gyrus |
| 13 | Fusiform_L (aal) |
| 9 | brodmann area 37 |
| 9 | brodmann area 37 |
| 8 | Limbic Lobe |
| 6 | Cerebelum_Crus1_L (aal) |
| 6 | Insula_R (aal) |
| 6 | Uncus |
| 5 | Limbic Lobe |
| 5 | Parahippocampa Gyrus |
| 5 | ParaHippocampal_R (aal) |
| 5 | Temporal_Inf_L (aal) |
| 4 | brodmann area 19 |
| 4 | Frontal_Inf_Orb_R (aal) |
| 4 | Insula |
| 4 | Sub-lobar |
| 3 | brodmann area 22 |
| 3 | brodmann area 28 |
| 3 | Inferior Frontal Gyrus |
| 3 | Inferior Temporal Gyrus |
| 3 | Temporal_Inf_L (aal) |
| 2 | Amygdala_R (aal) |
| 2 | brodmann area 13 |
| 2 | brodmann area 36 |
| 2 | brodmann area 47 |
| 2 | Cerebro-Spinal Fluid |
| 2 | Lateral Ventricle |
| 2 | Parahippocampa Gyrus |
| 2 | Sub-lobar |
| 2 | Superior Temporal Gyrus |
| 1 | Amygdala |
| 1 | brodmann area 20 |
| 1 | brodmann area 34 |
| 1 | Cerebellum Anterior Lobe |
| 1 | Culmen |
| 1 | Left Cerebellum |
| 1 | Olfactory_R (aal) |
| 1 | ParaHippocampal_L (aal) |
| 1 | Temporal_Pole_Sup_R (aal) |

**Table S3.**
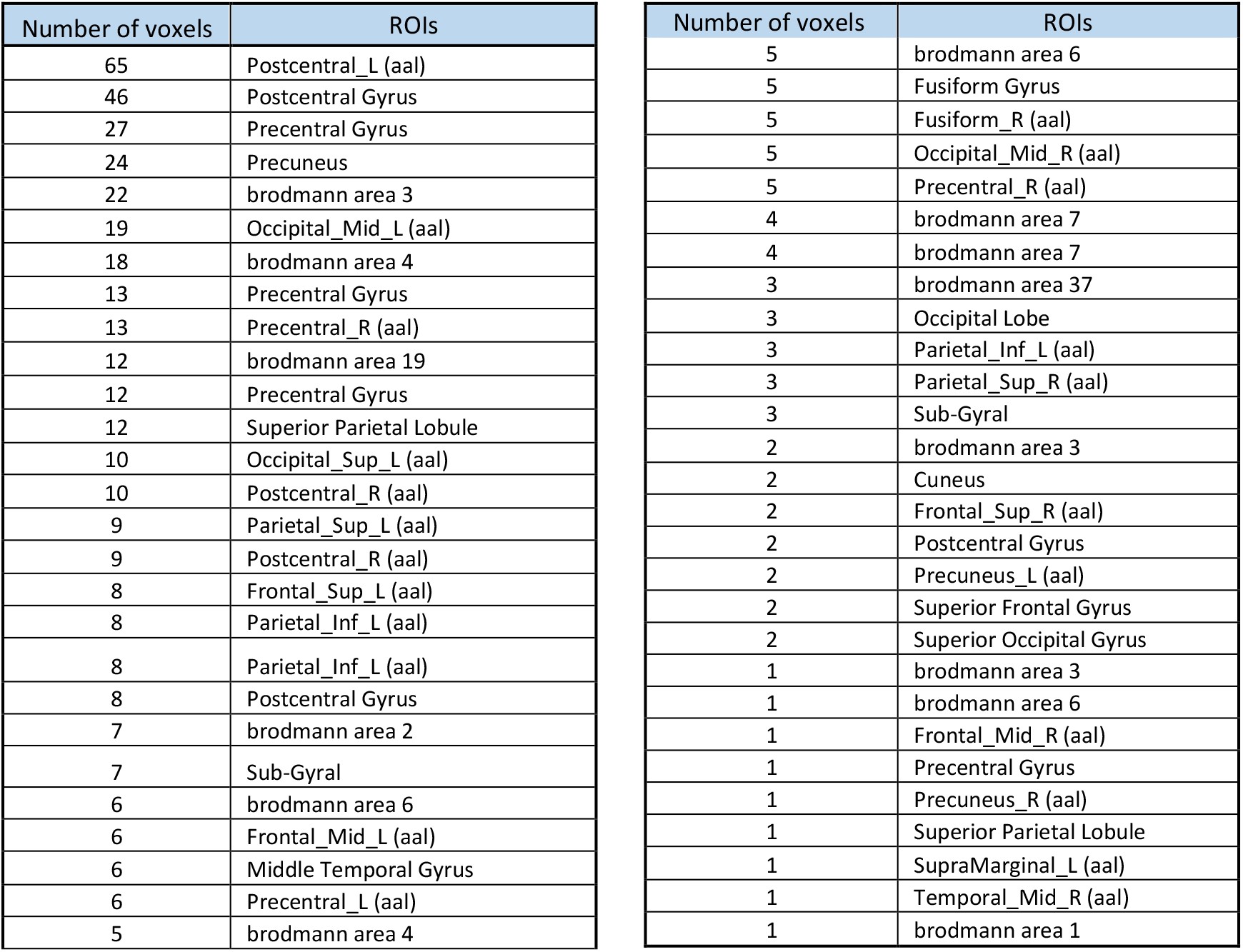
The number of voxels in each cluster that are activated during the choice task in Subject04. Subject04 had the lowest score in Novelty Seeking of TCI test. The contrast was defined as (self-difficult - other conditions) and the p-value threshold was set at (P<0.05).

**Table S4.**
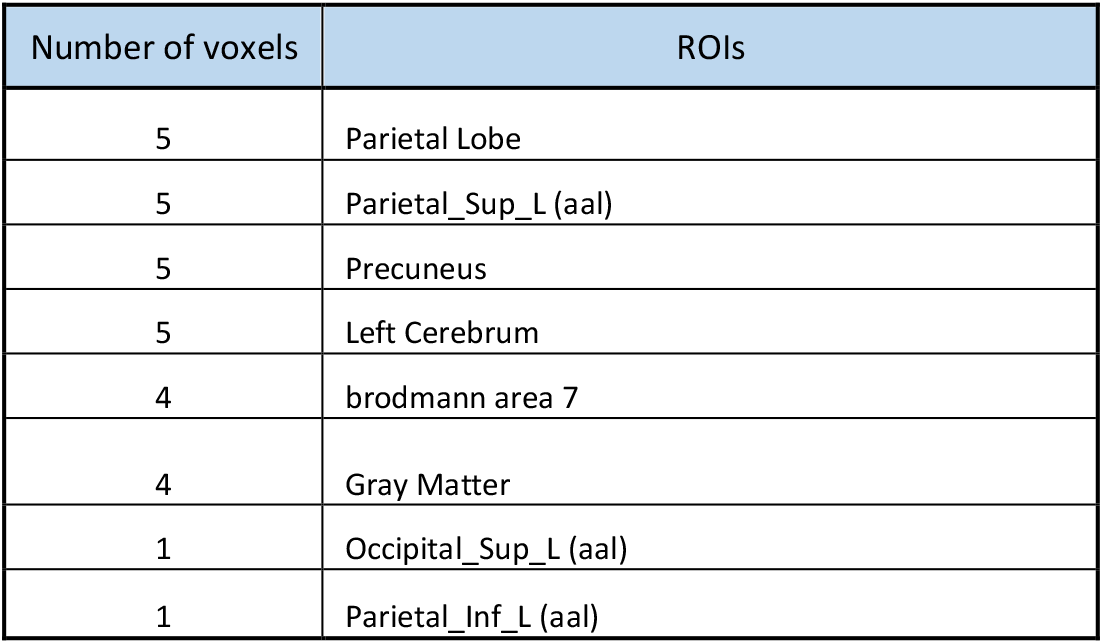
The number of voxels in each cluster that are activated during the choice task in Subject05. Subject05 had the highest score in Reward Dependence of TCI test. The contrast was defined as (self-difficult - other conditions) and the p-value threshold was set at (P<0.05).

| Number of voxels | ROIs |
| --- | --- |
| 5 | Parietal Lobe |
| 5 | Parietal_Sup_L (aal) |
| 5 | Precuneus |
| 5 | Left Cerebrum |
| 4 | brodmann area 7 |
| 4 | Gray Matter |
| 1 | Occipital_Sup_L (aal) |
| 1 | Parietal_Inf_L (aal) |

**Table S5.** The number of voxels in each cluster that are activated during the choice task in Subject07. Subject07 had the highest score in Harm Avoidance and the lowest score in Persistence of TCI test. The contrast was defined as (self-difficult - other conditions) and the p-value threshold was set at (P<0.05).

| Number of voxels | ROIs |
| --- | --- |
| 8 | Right Cerebrum |
| 8 | SupraMarginal_R (aal) |
| 7 | Supramarginal Gyrus |
| 6 | brodmann area 40 |
| 6 | Gray Matter |
| 5 | brodmann area 40 |
| 5 | Gray Matter |
| 4 | Frontal Lobe |
| 4 | Medial Frontal Gyrus |
| 3 | Gray Matter |
| 2 | brodmann area 10 |
| 1 | Anterior Cingulate |
| 1 | brodmann area 32 |
| 1 | Limbic Lobe |
| 1 | SupraMarginal_L (aal) |

**Table S6.** The number of voxels in each cluster that are activated during the choice task in Subject09. Subject09 had the highest score in Novelty Seeking of TCI test. The contrast was defined as (self-difficult - other conditions) and the p-value threshold was set at (P<0.05).

| Number of voxels | ROIs |
| --- | --- |
| 112 | Frontal Lobe |
| 48 | Precentral Gyrus |
| 43 | Precentral_L (aal) |
| 50 | Parietal Lobe |
| 31 | Postcentral_L (aal) |
| 26 | Postcentral Gyrus |
| 22 | Paracentral_Lobule_L (aal) |
| 21 | brodmann area 6 |
| 17 | Sub-Gyral |
| 12 | Parietal_Inf_L (aal) |
| 11 | Postcentral Gyrus |
| 9 | brodmann area 5 |
| 9 | Sub-lobar |
| 9 | Middle Frontal Gyrus |
| 15 | brodmann area 2 |
| 9 | Cingulate Gyrus |
| 9 | Limbic Lobe |
| 7 | Frontal_Sup_R (aal) |
| 7 | Inferior Parietal Lobule |
| 7 | Parietal_Sup_L (aal) |
| 12 | Sub-Gyral |
| 7 | Supp_Motor_Area_R (aal) |
| 6 | Cerebro-Spinal Fluid |
| 6 | Lateral Ventricle |
| 6 | Postcentral_L (aal) |
| 5 | brodmann area 4 |
| 5 | brodmann area 40 |
| 5 | Frontal_Sup_R (aal) |
| 5 | Superior Frontal Gyrus |
| 9 | Medial Frontal Gyrus |
| 9 | Middle Frontal Gyrus |
| 6 | Sub-Gyral |
| 3 | brodmann area 3 |
| 3 | brodmann area 40 |
| 3 | Extra-Nuclear |
| 3 | Frontal_Sup_L (aal) |
| 3 | Sub-lobar |
| 3 | Superior Frontal Gyrus |
| 3 | Supp_Motor_Area_R (aal) |
| 2 | brodmann area 6 |
| 2 | Corpus Callosum |
| 2 | Extra-Nuclear |
| 2 | Precentral_R (aal) |
| 1 | brodmann area 24 |
| 1 | Caudate |
| 1 | Caudate Tail |
| 1 | Paracentral Lobule |

**Table S7.** The number of voxels in each cluster that are activated during the choice task in Subject11. Subject11 had the highest score in Persistence of TCI test. The contrast was defined as (self-difficult - other conditions) and the p-value threshold was set at (P<0.05).

| Number of voxels | ROIs |
| --- | --- |
| 10 | Precuneus_L (aal) |
| 7 | Frontal Lobe |
| 7 | Paracentral Lobule |
| 7 | Parietal Lobe |
| 6 | Occipital_Mid_L (aal) |
| 6 | Postcentral Gyrus |
| 5 | Cerebellum Posterior Lobe |
| 5 | Right Cerebellum |
| 4 | Cerebellar Tonsil |
| 3 | Middle Occipital Gyrus |
| 2 | Cerebelum_7b_R (aal) |
| 2 | Cerebelum_8_R (aal) |
| 2 | Paracentral_Lobule_R (aal) |
| 2 | Right Cerebrum |
| 1 | brodmann area 4 |
| 1 | brodmann area 5 |
| 1 | brodmann area 6 |
| 1 | Cerebelum_Crus2_R (aal) |
| 1 | Inferior Semi-Lunar Lobule |
| 1 | Sub-Gyral |
| 1 | brodmann area 18 |

**Table S8.**
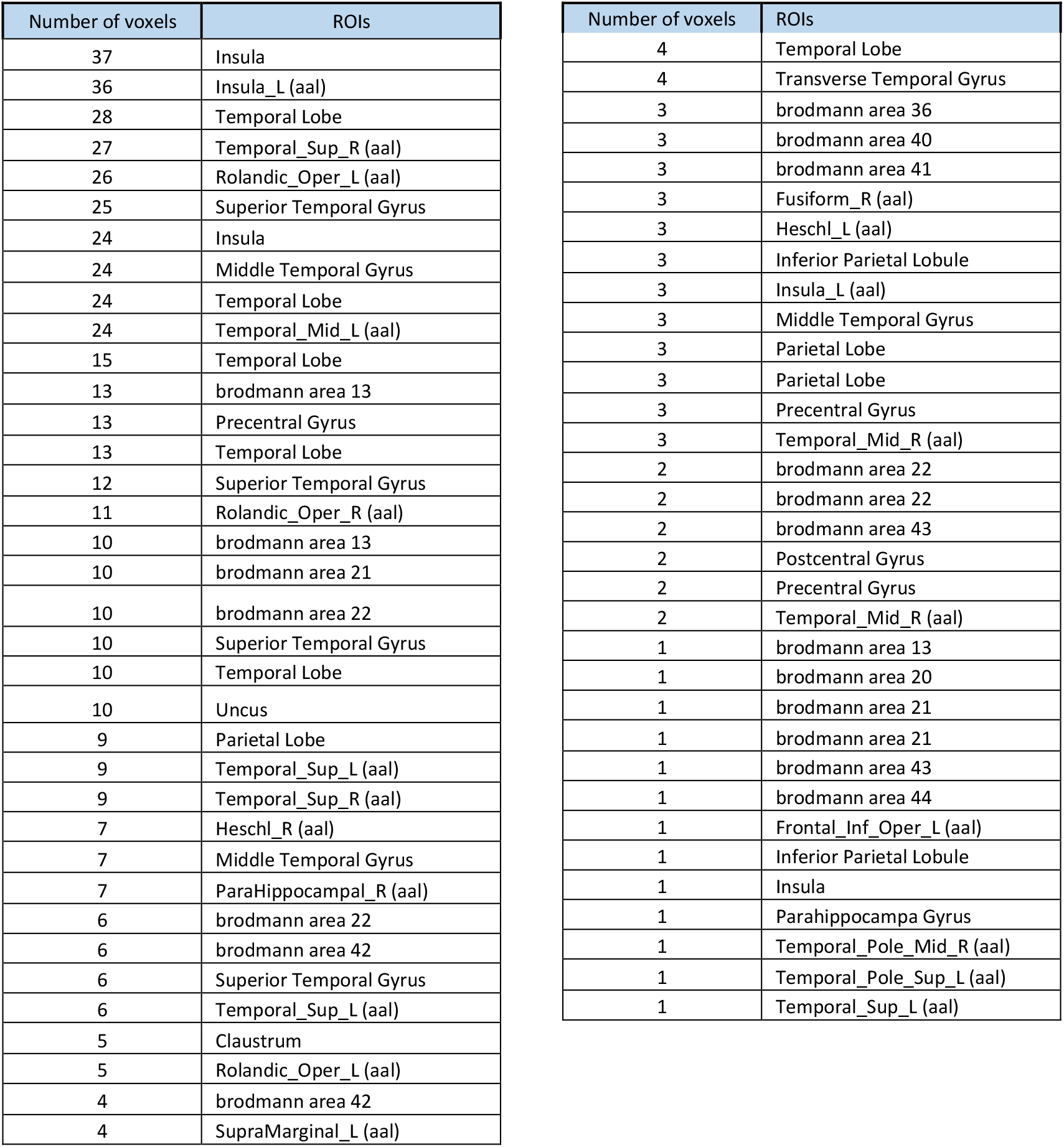
The number of voxels in each cluster that are activated during the choice task in Subject13. Subject13 had the lowest score in Harm Avoidance of TCI test. The contrast was defined as (self-difficult - other conditions) and the p-value threshold was set at (P<0.05).

**Table S9.** Demographic characteristics of the participants who did the behavioral test.

| Characteristics | class | Frequency | Frequency (%) |
| --- | --- | --- | --- |
| Gender | Male | 112 | 48 |
|  | Female | 122 | 52 |
| Age (Years) | <20 | 16 | 7 |
|  | 20-30 | 80 | 34 |
|  | 30-40 | 82 | 35 |
|  | 40-50 | 40 | 17 |
|  | >50 | 16 | 7 |
| Educational qualifications | Highschool diploma | 56 | 24 |
|  | Bachelor's degree | 82 | 35 |
|  | Master's and higher degree | 96 | 41 |

